# Mucin-derived *O*-glycans supplemented to diet mitigate diverse microbiota perturbations

**DOI:** 10.1101/2020.06.09.140897

**Authors:** K. M. Pruss, A. Marcobal, A.M. Southwick, D. Dahan, S.A. Smits, J.A. Ferreyra, S.K. Higginbottom, E.D. Sonnenburg, P.C. Kashyap, B. Choudhury, L. Bode, J. L. Sonnenburg

## Abstract

Microbiota-accessible carbohydrates (MACs) are powerful modulators of microbiota composition and function. These substrates are often derived from diet, such as complex polysaccharides from plants or human milk oligosaccharides (HMOs) during breastfeeding. Host-derived mucus glycans on gut-secreted mucin proteins may serve as a continuous endogenous source of MACs for resident microbes; here we investigate the potential role of purified, orally-administered mucus glycans in maintaining a healthy microbial community. In this study, we liberated and purified *O*-linked glycans from porcine gastric mucin and assessed their efficacy in shaping the recovery of a perturbed microbiota in a mouse model. We found that porcine mucin glycans (PMGs) and HMOs enrich for taxonomically similar resident microbes. We demonstrate that PMGs aid recovery of the microbiota after antibiotic treatment, suppress *C. difficile* abundance, delay the onset of diet-induced obesity, and increase relative abundance of resident *Akkermansia muciniphila. In silico* analysis revealed that genes associated with mucus utilization are abundant and diverse in prevalent gut commensals and rare in enteric pathogens, consistent with these glycan-degrading capabilities being selected for during host development and throughout evolution of the host-microbe relationship. Importantly, we identify mucus glycans as a novel class of prebiotic compounds that can be used to mitigate perturbations to the microbiota and provide benefits to host physiology.

## Introduction

The luminal surface of the gastrointestinal tract is covered by a viscous mucus layer, which serves as the primary interface at which the host interacts with a dense microbial community. Secreted by host goblet cells, mucus is largely composed of highly glycosylated mucin proteins. The gel-like mesh of mucins provides both a barrier to shield the host from direct interaction with microbes, preventing inflammation[1], but also provides an energy-rich substrate for the microorganisms that reside in the gut[2, 3]. Degradation of the diverse chemical linkages within endogenous glycans requires a specialized set of glycoside hydrolases, reflected in the genetic composition of the gut microbiota. Notably, species prevalent in the human gut microbiota often possess broad glycan-degrading capabilities while “specialist” species may have narrower glycan-degrading potential[4].

The structural features of intestinal mucus glycans are strikingly similar to those of human milk oligosaccharides (HMOs)[5]. Mucin glycans are built upon an *N-*acetylgalactosamine that is *O*-linked to serine and threonine residues of the mucin protein, while HMOs are built upon a lactose core structure universally present at the reducing end of these glycans[6]. In both mucin glycans and HMOs, the priming carbohydrate structure is extended with galactose-*N-*acetylglucosamine disaccharides and chains often terminate with fucose or sialic acid residues[7].

HMOs are the third most abundant compound in breast milk, after lactose and fat[8]. Colostrum is particularly rich in HMOs, the concentration ranging from 20-25 g/L in the first milk produced and decreasing to 5-20 g/L in more mature milk[9–11]. Numerous and diverse beneficial effects have been attributed to HMOs in breast milk during infant development[12–20].

Recently, extensive research has focused on the potential prebiotic role of HMOs. HMOs are indigestible by humans and are degraded throughout the gastrointestinal tract of breast-fed infants[21], becoming the primary microbiota-accessible carbohydrates available in the newborn diet. Species belonging to the genera *Bifidobacteria* and *Bacteroides* are optimal HMO-consumers: *Bifidobacterium longum* subsp. *infantis* and *Bifidobacterium bifidum* encode in their genomes clusters of genes dedicated to HMO utilization[22–24]. *Bacteroides thetaiotaomicron*, described as a “generalist-glycan-consumer” due to the numerous glycoside hydrolases encoded in its genome, grows efficiently on HMOs *in vitro*[5, 25]. *In vivo* studies with gnotobiotic mice revealed that lacto-*N*-neotetraose, one of the most abundant oligosaccharides in HMOs, provides an advantage to *B. infantis* over *B. thetaiotaomicron* in gut colonization[5]. However, the *in vivo* effect of the structurally diverse HMO pool in microbiota composition remains under-explored.

In the infant gut, MAC-consuming bacteria that reach the intestinal tract must rely primarily on HMOs or mucus as growth substrates. Interestingly, transcriptional data demonstrates that HMO utilization in *Bacteroides* and *Bifidobacterium* relies on pathways that also play a role in mucus utilization[5]. As HMOs have been described to confer numerous benefits to infants and similarities between microbial metabolism of HMOs and mucin glycans has been shown, we investigated the extent to which mucin glycans are able to confer benefits to the microbiota and host. We hypothesized that targeting convergent glycan-utilization pathways in the adult gut community with exogenously-administered porcine mucin glycans (PMGs) could mitigate perturbations to the microbiota.

Empirical studies of the human microbiota can be performed in a controlled environment by colonizing germ free (GF) mice with specific strains of bacteria (gnotobiotic) or a complete microbial community from human feces (humanized). The humanized mouse model recapitulates the vast majority of human microbiota compositional and functional features[26, 27]. In this work, we use both gnotobiotic and humanized mice to establish that the gut microbiota efficiently consumes HMOs. Furthermore, we demonstrate that a complex mix of glycans isolated from porcine mucin recreates some of the effect of HMOs on the gut microbiota and mitigates the negative effects of various community perturbations including antibiotic treatment, pathogen invasion, and a high-fat diet.

## Results

### HMOs are consumed by members of the microbiota, conferring a growth advantage to *Bifidobacterium* over *Bacteroides in vivo*

We and others have previously reported the ability of both *Bacteroides* and *Bifidobacterium* to utilize select HMOs *in vitro*[5, 28]; as such, we wished to determine whether complex HMOs isolated from human donors favored *Bifidobacterium* or *Bacteroides* within the context of the gut environment. 6-week old GF mice were bi-colonized with two HMO-utilizing taxa common to the infant gut: *Bacteroides thetaiotaomicron* (*Bt*) and *Bifidobacterium longum* subsp *infantis* (*B. infantis*). Mice were fed a MAC-deficient (MD) diet supplemented with HMOs (1% in water, chosen to approximate the mass of HMOs consumed by human newborns, adjusted for body weight, see Methods) for one week. Feces and cecal contents were collected and milk glycans were measured by HPLC from all samples. In bi-colonized mice, no HMOs were detectable, whereas GF mice samples revealed a high concentration of milk glycans in both the fecal (**Fig 1A)** and cecal samples **(Fig S1A, B**), demonstrating that HMO-utilizing members of the commensal microbiota deplete HMOs within the host large intestine.

**Figure 1.**
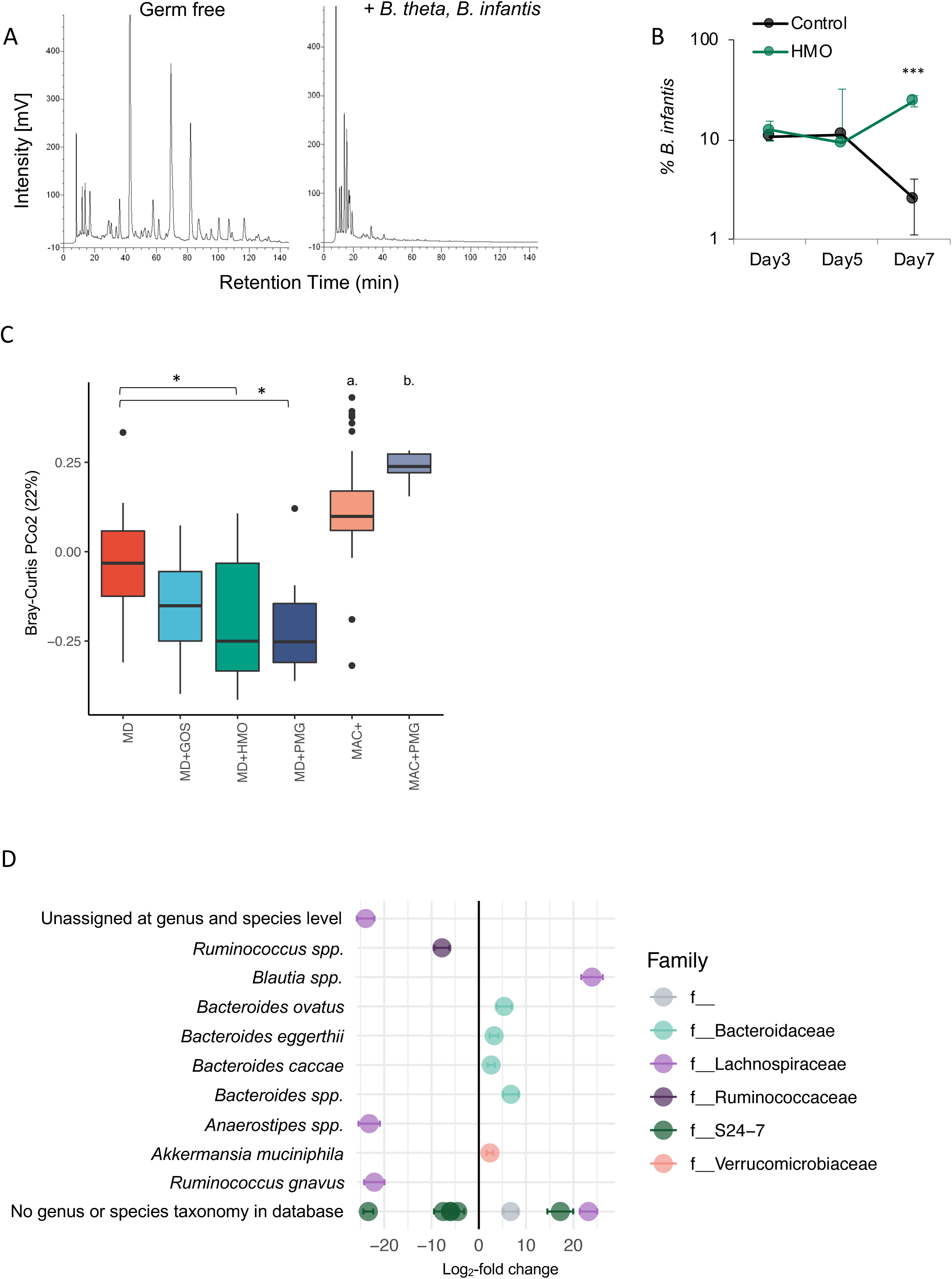
Human milk oligosaccharides (HMOs) are consumed by and shape the commensal microbiota. **A)** Germ-free and mice bi-colonized with *Bt* and *B. infantis* were fed MD diet supplemented with 1% HMOs (w/v in water). HPLC-FL based chromatograms of glycan content in stool samples at day 7 reveal degradation of HMOs *in vivo*. **B)** HMO supplementation provides a competitive advantage to *B. infantis* over *Bt* in bi-colonized mice (n=4 mice/group, mean +/- SEM, *** = *P* < 0.001). Abundance of *Bt* and *B. infantis* from feces was determined with CFU dilution plating. **C)** Bray-Curtis dissimilarity metric indicates significant differences to community composition between mice on MD diet and MD diet supplemented with 1% PMGs, or HMOs. MD diet alone is not significantly different from MD diet supplemented with 1% GOS. (Data were combined across sampling time-points: n=12 MD, n=19 MD+GOS, n=21 MD+HMOs, n=11 MD+PMG, n=61 MAC^+^, n=10 MAC^+^+PMG. a. * MAC^+^ vs. MD; **** MAC+ vs. MD+GOS, MD+HMOs, and MD+PMG. b. *** MAC^+^+PMG vs. MD; **** MAC++PMG vs. MD+GOS, MD+HMOs, and MD+PMG. *** *P* < 0.001, **** *P* < 0.0001, pairwise t-tests with Bonferroni multiple hypothesis correction). **D)** Individual ASVs with significantly different abundance due to HMO supplementation (positive log_2_-fold change) versus MD diet alone (negative log_2_-fold change, mean +/- SEM, adjusted *P*-value < 0.01, parametric Wald test). Highest resolution taxonomic assignment indicated to the left.

*In vitro, Bt* grows better on mucus glycans than *B. infantis*, whereas *B. infantis* grows better on HMOs than *Bt*[5]. After 1 week, HMO supplementation resulted in an expansion in the population of *B. infantis* relative to *Bt* compared to mice on regular water (24.3 ± 3.12 % versus 2.5 ± 1.45 % on day 7; *P* < 0.01, n=4 mice) (**Fig 1B)**. These results confirm that HMOs provide a selective advantage to *B. infantis* over *Bt in vivo*. Therefore, although HMOs are utilized by generalist glycan degraders, they can provide a competitive advantage to microbes that specialize in HMO utilization.

### HMOs shape the composition of the gut microbiota

To address the extent to which HMOs shape a complex microbial community, we administered purified HMOs to ex-GF mice colonized for 6 weeks with a human microbiota (humanized). Compared to either a standard mouse diet rich in MACs (MAC^+^) diet or MD diet alone, purified HMOs engender a distinct microbial community (**Fig 1C)**. HMO supplementation to MD diet led to the significant enrichment of the known mucin glycan degraders *Bacteroides caccae* and *Akkermansia muciniphila*[25], as well as polysaccharide generalists *Bacteroides ovatus* and *Bacteroides eggerthii* (**Fig 1D**).

Purified HMOs are structurally diverse, comprised of many molecules with different chemical properties[6]. An important question is whether glycans with slight structural differences within HMO mixtures can differentially impact the community. To investigate whether structural nuance in exogenous glycans led to changes in the composition of the gut microbiota, we employed two of the most abundant glycans found in HMOs and administered them in pure form to humanized mice. Lacto-*N-*tetraose (LNT, Galβ1-4GlcNAcβ1-3Galβ1-4Glc) and Lacto-*N*-neotetraose (LNnT, Galβ1-3GlcNAcβ1-3Galβ1-4Glc) differ only in the placement of a linkage between galactose and *N*-acetylglucosamine (**Fig S1C)**. Germ-free mice were humanized and then switched to MD supplemented with LNT or LNnT (1% w/v in water) or maintained on MAC^+^ with plain water.

Bray-Curtis dissimilarity metric reveals clear separation of microbiota composition between mice consuming a MAC^+^ diet from MD supplemented with either of the two HMOs. Moderate but significant separation in composition was also observed between LNT versus LNnT supplementation in the MD diet (**Fig S1D**). The switch from MAC^+^ to MD background diet drives high-level structural changes to the composition of the humanized gut microbiota (**Fig S1E**). However, nuanced higher-resolution taxonomic changes occur corresponding to supplementation of one of two isomeric tetrasaccharides that differ in a single glycosidic linkage: nine taxa were significantly different at adjusted *P* value < 0.01 between LNT and LNnT supplementation (parametric Wald test[29], data not shown), including four ASVs belonging to the Bacteroidales family S24-7, *Coprococcus spp*. and *Sutterella spp*., two unidentified Erysipelotrichaceae (enriched with LNnT), and one unidentified *Blautia* strain (LNT). Accordingly, a Random Forests classifier predicts the three diets with 87.5% accuracy (out-of-bag estimate of error, with 30% class error for LNnT, 10% for LNT, and 0% for SD. Leave-one-out cross-validation results in accuracy of 94%), supporting the subtle but significant differences imparted onto the gut community by two synthetic oligosaccharides.

### Structural analysis reveals similarities between PMGs and HMOs

HMOs are structurally similar to mucin glycans. We modified a described protocol[30] for purifying neutral *O*-glycans from commercially available porcine mucin using reductive ß-elimination followed by anion exchange chromatography, and characterized the purified material. Hydrolysis with TFA and HPAEC-PAD analysis revealed that PMGs are composed of units of glucosamine, galactosamine, galactose, glucose, fucose and mannose, consistent with previous reports of mucin glycan composition ([31], **Fig 2A)**. The unexpected presence of mannose, a monosaccharide not commonly found in *O*-linked glycans is likely due to a small amount of liberated *N-*glycan during purification. MALDI-TOF mass spectrometry identified thirteen major glycans (**Fig 2B**), for which structures were predicted with Glycobench software (see Materials and Methods) based on m/z values, and three of thirteen were confirmed by MS/MS fragmentation pattern (**Fig 2C, Fig S2)**. Structures were inferred for the remaining ten masses based on previous structural work on mucin glycans **(Fig 2D)**. Sialic acid content quantification from purified PMGs revealed an absence of *N*-acetylneuraminic acid and *N*-glycolylneuraminic acid (data not shown) consistent with anion chromatographic depletion of the negatively charged glycan fractions. Our analysis affirms the similarity between PMGs and HMOs both in terms of structure and constitutive components.

**Figure 2.**
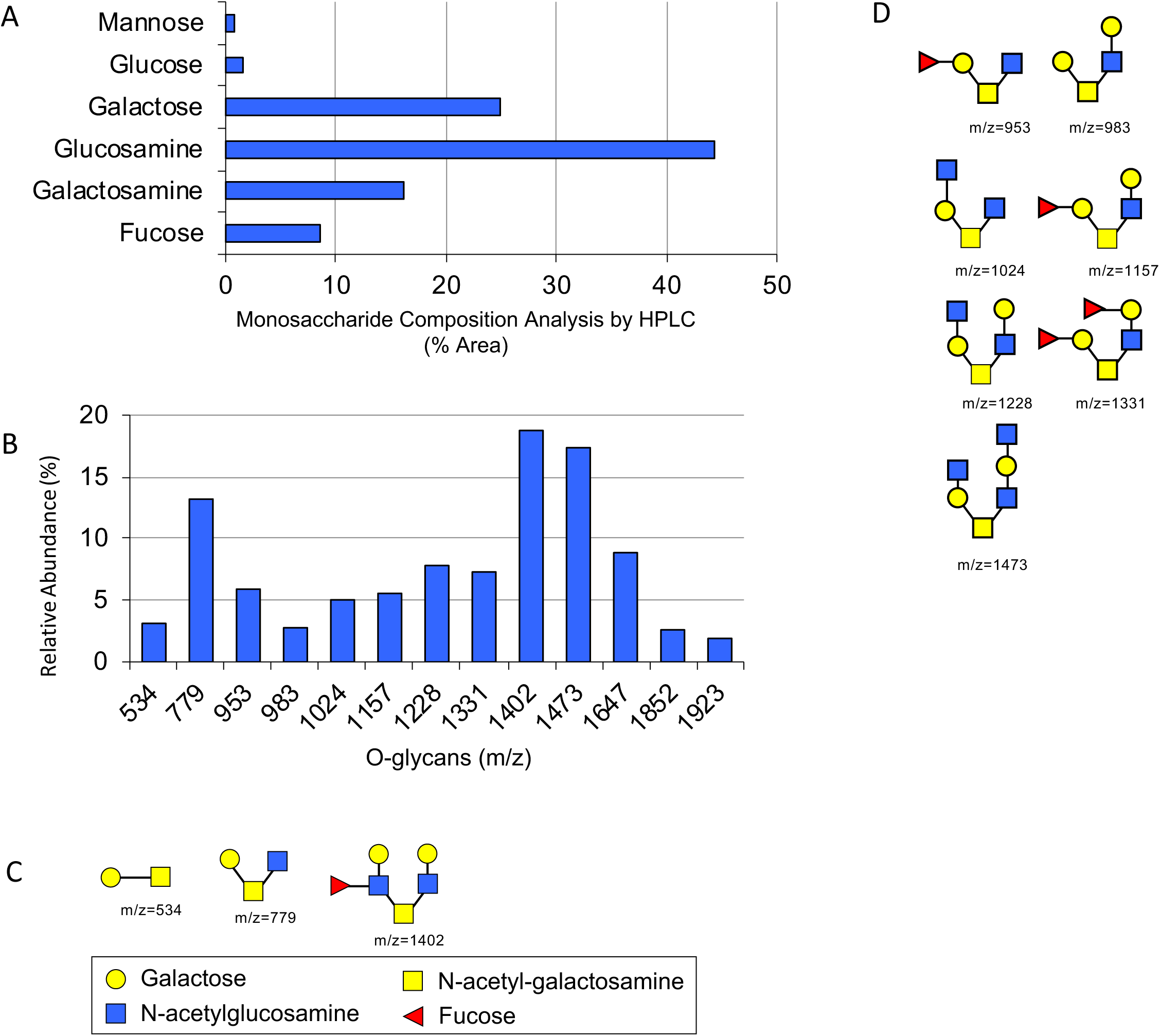
Structural analysis of porcine mucin glycans (PMGs). **A)** Six detectable monosaccharides were identified after total acid hydrolysis of PMGs, with four monosaccharides associated with mucin glycans dominating. Amino sugars are the N-acetyl forms. **B)** 13 abundant *O*-linked glycans were detected in purified PMGs using MALDI-TOF mass spectrometry. **C**,**D)** Structures of the *O-*glycans quantified in **(B)** were predicted with GlycoWork Bench. Three of the identified PMGs **(C)** were validated with MS/MS.

### PMGs and HMOs drive similar high-level microbial community changes

Given the structural similarities between PMGs and HMOs, we sought to determine how exogenously administered PMGs would affect the composition of the gut microbiota. We administered pools of PMGs (1% in water, to MD or MAC^+^ diets) or a common synthetic prebiotic, galacto-oligosaccharide (GOS, 1% in water to MD) and compared microbiota composition between these conditions and the HMO supplementation experiment described above. Similar high-level changes in the community were observed when HMOs, PMGs or GOS were supplemented to a diet deficient in plant polysaccharides, indicating taxonomic changes at the family level are driven primarily by background diet (**Fig S3A)**. A switch from MAC+ background diet to MD led to a reduction in alpha diversity, which was not restored by administration of any exogenous glycans tested (**Fig S3B)**. However, administration of 1% PMGs to MAC^+^ background diet led to a significant increase in alpha diversity (**Fig S3B**). To support this finding, we sought to determine whether a lower concentration of PMG supplementation would yield the same enrichment, and indeed found that 0.3% PMGs in water supplemented to a MAC^+^ diet led to a similar increase in alpha diversity (**Fig S3C**).

While the first principal component of Bray-Curtis dissimilarity is driven by background diet (MAC^+^ versus MD), the second principal component reveals significant separation between communities supplemented with PMGs or HMOs from MD alone, whereas GOS supplementation is not significantly different from MD (**Fig 1C**). The gap statistic predicts 5 clusters amongst the 6 diets (MAC^+^ +/- 1% PMG; MD +/- 1% PMG, HMO or GOS) (canonical correspondence analysis, method = firstSEmax[32]), providing further support for nuanced divergence of microbial communities in response to exogenous glycan administration. As seen with the administration of pure LNT and LNnT, administration of PMGs or HMOs to MD diet led to significant changes to the community at lower taxonomic levels. Both HMO (**Fig 1D**) and PMG supplementation (**Fig S3D**) led to the significant enrichment of the mucin-degrader *B. caccae* and an unidentified member of the *Blautia* genus. PMG supplementation to MD led to a significant increase in *Bacteroides eggerthii*, whereas *B. fragilis* and *B. ovatus* were enriched with PMG supplementation to MAC^+^ diet (**Fig S3E**).

### PMGs accelerate recovery from antibiotic perturbation

We next pursued investigation of purified PMGs to mitigate microbiota disturbance. Humanized mice were switched to MD diet and treated with 1 mg clindamycin concurrent with 1% PMG supplementation in water. When compared with no supplementation, PMG led to faster recovery of alpha diversity (**Fig 3A)** and an accelerated trajectory to the baseline community microbiota as measured by comparison of UniFrac distance to pre-antibiotic timepoints (**Fig 3B)**. Furthermore, mice treated with PMGs exhibit a reduced bloom of Proteobacteria, a hallmark of post-antibiotic oxygenation and inflammation in the gut (**Fig 3C, S4A**, [33, 34]). Additionally, PMG supplementation with antibiotics leads to a faster recovery of the relative abundance of *A. muciniphila* (**Fig S4B)**. These data suggest that exogenous glycans could be simultaneously administered with a course of antibiotics to aid in recovery to the microbiota.

**Figure 3.**
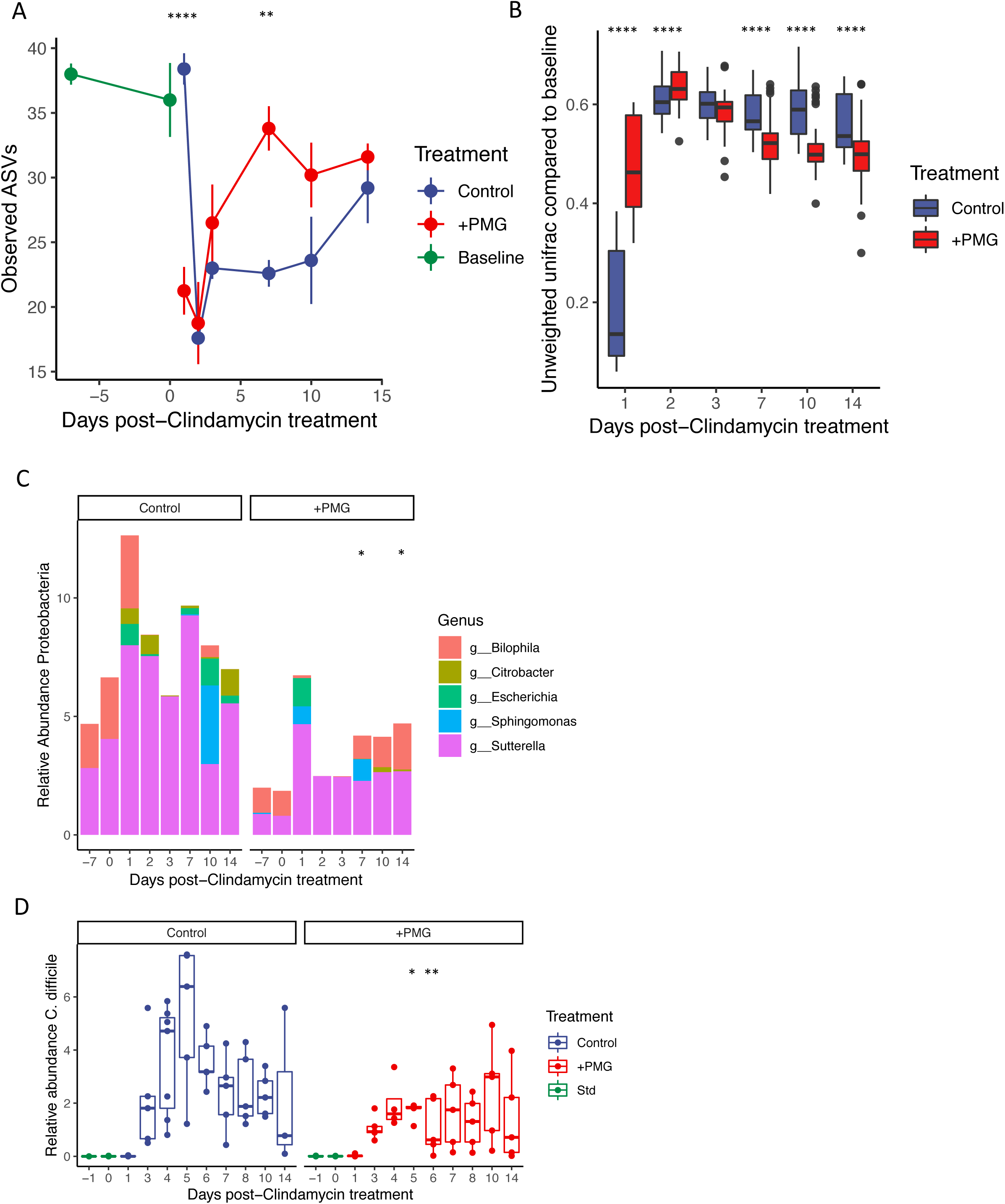
Treatment with PMGs leads to accelerated post-antibiotic recovery. **A)** MD diet supplemented with 1% PMGs (+PMG) leads to accelerated recovery of alpha diversity compared to MD diet alone (Control, mean +/- SEM shown, ** *P* < 0.01, **** *P* < 0.0001, pairwise t-tests with Bonferroni correction for post-antibiotic treatment timepoints) Baseline alpha diversity measurements are during MAC^+^ diet prior to clindamycin treatment. **B)** Unweighted UniFrac distance (compared to pre-antibiotic MAC^+^ baseline) reveals that PMG supplementation to MD diet (+PMG) leads to a faster trajectory back to baseline community than MD diet alone (Control, **** *P* < 0.0001, pairwise t-tests with Bonferroni correction). **C)** Mean relative abundance (%) of the phylum Proteobacteria is suppressed in mice treated with 1% PMGs compared to the MD control. Bars are colored by genus. Total Proteobacteria abundance is significantly higher days 7 and 14 post-antibiotic treatment in the control group. (* *P* < 0.05, pairwise t-tests with Bonferroni correction). **D)** Relative abundance of *Cd* is suppressed with PMG supplementation compared to MD diet alone (* *P* < 0.05, ** *P* < 0.01, pairwise t-tests with Bonferroni correction).

### PMGs suppress *C. difficile* abundance

As PMGs augmented community recovery post-antibiotics, we sought to determine whether exogenous PMGs would affect host susceptibility to an antibiotic-associated pathogen, *Clostridium difficile* (*Cd*). *Cd* depends on disturbance to the microbiota to cause disease [35, 36] and thus represented a suitable target to determine whether exogenous mucus glycans could protect the host from infection.

1% PMGs were supplemented to MAC^+^ or MD diet, concomitant with antibiotic treatment (1 mg clindamycin). Mice were gavaged 24 hours later with 200 µl saturated overnight culture wild-type *Cd* 630 (Day 1). Administration of PMGs to mice on a background diet devoid of complex polysaccharides significantly reduced the *Cd* burden as measured by 16S rRNA reads (**Fig 3D)** and selective plating (**Fig S5A)**. PMGs supplemented to MAC^+^ diet did not affect *Cd* abundance (**Fig S5B)** or histopathology (**Fig S5C**). Whether the inhibitory effect of PMGs towards *Cd* occurs via reshaping the microbial community in a way such that *Cd* suffers a competitive disadvantage, directly inhibiting *Cd* growth or toxin activity[37], or altering host immune signaling directly or indirectly via the microbiota remains to be determined.

### PMGs attenuate host weight gain due to High Fat Diet

As *A. muciniphila* has been associated with diet-induced obesity models previously [38–40] and appears to be manipulable with administration of PMGs, we were interested in whether administration of PMGs could attenuate the effects of a high-fat diet (HFD) on host physiology. Three groups of age- and sex-matched mice were fed either a standard diet, HFD (60% fat and 20% carbohydrates), or HFD supplemented with 1% PMGs continuously in drinking water and their weight monitored over three weeks. HFD induced significant weight gain compared to MAC^+^-diet fed controls, and PMG supplementation to HFD significantly reduced host weight gain (**Fig 4A)** and development of adipose tissue (**Fig 4B)**, despite no differences between the groups in the amount of food consumed. No differences in glucose tolerance were observed between HFD-fed mice and PMG supplementation to HFD (data not shown).

**Figure 4.**
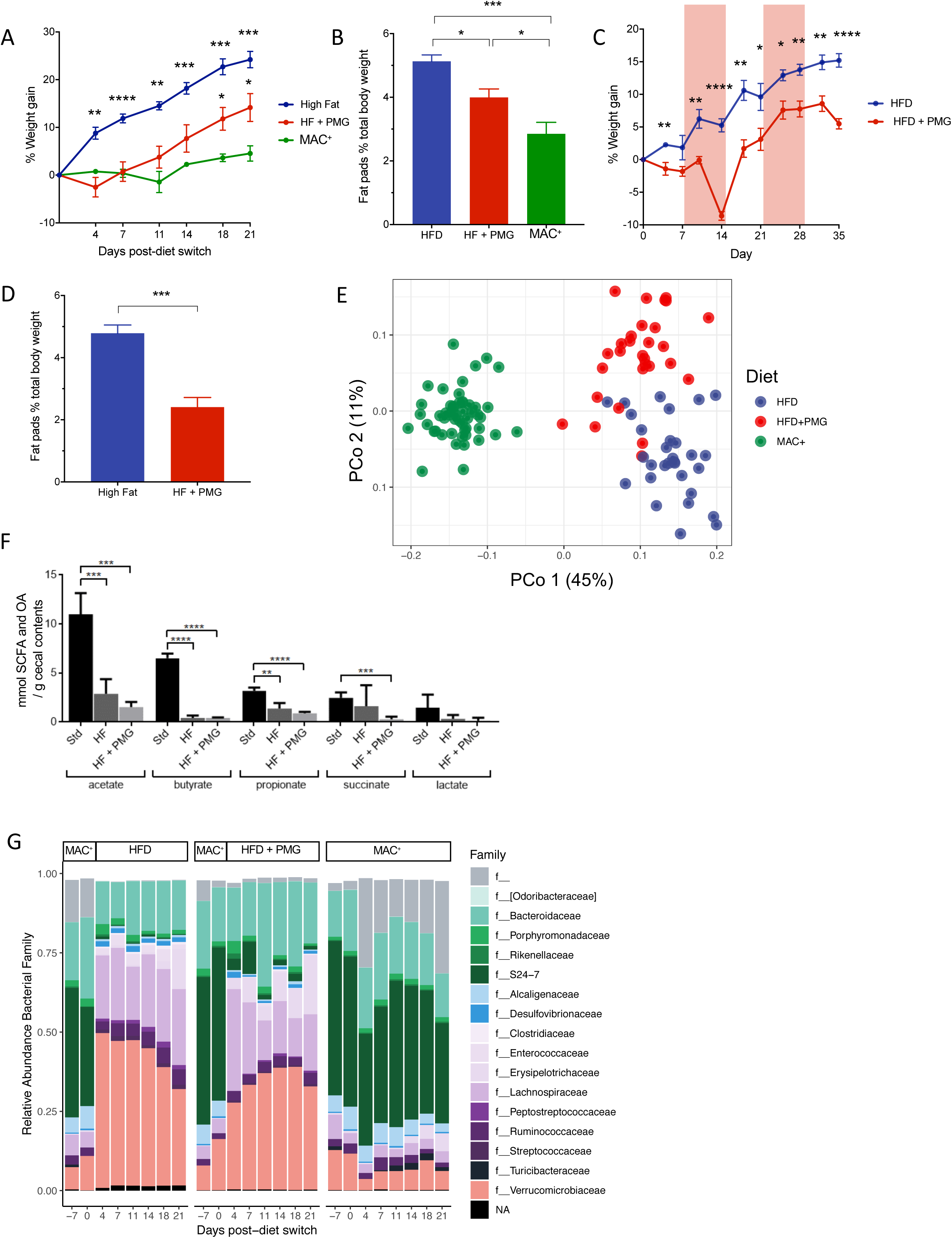
PMGs mitigate the effect of high-fat diet on host physiology and the gut microbiota. **A)** Weight gain in individual mice fed HFD (n=5 mice, blue), HFD supplemented continuously with 1% PMGs (n=5, red), or maintained on MAC^+^ diet (n=4, red). Weight was compared to baseline day 0 prior to diet switch (mean +/- SEM, repeated measures two-way ANOVA with Dunnett’s multiple comparison’s test, * indicates significance versus MAC^+^). **B)** Fat pads as percent total body mass at day 21 (mean +/- SEM, ANOVA). **C)** PMGs administered for 7-day durations in water (1% w/v) are sufficient to reduce host weight gain **(C)** and fat accumulation **(D)** due to HFD. **C)** Weight gain compared to day 0 (mean +/- SEM, n = 5 mice/group, multiple t-tests). Red boxes denote period of two one-week pulses of PMGs administered to HFD + PMG group. **D)** Fat pads as percent total body mass at day 35 (mean +/- SEM, n = 5 mice/group). **E)** Unweighted UniFrac reveals separation of the three diet groups; PMG supplementation to HFD leads to a unique microbial community from HFD alone. **F)** Diet-dependent decreases in cecal short-chain fatty acids were determined by GC-MS in cecal contents of mice fed MAC^+^, HFD, or HFD + 1% PMG. **G)** Changes in the top 100 most abundant taxa over time for mice maintained on MAC^+^, HFD, or HFD supplemented with 1% PMGs. f_ indicates that a strain is not assigned at the family level in the Greengenes database; NA indicates lack of taxonomic assignment at the family level. For A-D,F: * *P* < 0.05, ** *P* < 0.01, *** *P* < 0.001, **** *P* < 0.0001.

Exogenous PMG administration to HFD led to a microbial community distinct from that of MAC^+^ or HFD alone. Although HFD induces similar family-level taxonomic changes with or without PMG supplementation (**Fig 4G**), beta-diversity analysis reveals clear separation of the humanized microbial communities on the three diets, indicating strong diet-driven changes in community composition (**Fig 4E, Fig S6A**). Accordingly, Random Forests classifies microbiota samples into the three diets perfectly (HFD, HFD+PMG, MAC^+^), and regression on percent weight gain in individual mice from baseline perfectly predicts diet. Predictive features from the Random Forests models include many members of the family S24-7, genera *Ruminococcus* and *Eubacterium* (data not shown). Short-chain fatty acid (SCFA) and organic acid (OA) analysis demonstrates drastically reduced metabolic output of microbial communities on HFD with and without PMGs (**Fig 4F**), indicating that the phenotypic effect of PMGs during HFD is not mediated by normalization of SCFA production.

To determine whether a shorter duration of PMG administration could attenuate host weight gain during HFD, humanized mice were switched to HFD and dosed with 1% PMGs for seven days at 1 and 4 weeks after diet change. Again, exogenous PMG administration led to less weight gain and significant reduction of fat accumulation (**Fig 4C**,**D)**. Two one-week pulses of PMGs led to a distinct microbial community from HFD alone; gut microbiota composition of mice dosed with PMGs remains distinct from the HFD-only group even when PMGs are removed from water (**Fig S6C)**. Random Forests again perfectly classifies the microbial communities into the groups on HFD alone or HFD supplemented transiently with PMGs. HFD leads to dramatic changes to the relative abundance of bacterial families compared to MAC^+^ diet (**Fig S6B**). Several taxa are significantly enriched due to exogenous PMG administration on HFD background, including *A. muciniphila* (**Fig S6D**), *B. caccae*, and members of the Lachnospiraceae family, including *Dorea, Coprococcus, Coprobacillus, Clostridium hathewayi, Blautia*, and *Eubacterium dolichum* (Wald test, adjusted *P* value < 0.0005, **Fig S6E**), which were also many of the top predictive features of the Random Forests classification (**Table S1**). There are multiple potential mechanisms by which PMGs could lead to reduced weight gain in mice, which is an area of important follow-up investigation.

### Mucin glycan utilization gene clusters are abundant in prevalent commensals and rare in pathogens

Given the structural similarities between PMGs and HMOs and the capabilities of PMGs to mitigate community disturbance, we hypothesized that mucus glycan utilization has played a role in host and commensal microbiota co-evolution. To explore this hypothesis, we performed a broad *in silico* search for candidate mucin-degrading carbohydrate gene clusters (CGCs) within the genomes of 4,500 common human gut commensals in the HGM database[41]. A CGC was defined as the colocalization of at least two glycoside hydrolases (GH) that have been previously reported to act on mucin (**Table S2**), plus a transporter or transcription factor within the same genomic locus. The rationale for requiring two GHs per locus was to increase the stringency in identifying candidate mucin CGCs: while these GH families contain members that act on mucus glycan structures, some of these families may also contain members that do not target mucin glycans. Therefore, by requiring two candidate mucin-degrading GHs we sought to reduce the instances of false positives, accepting that some mucin-degrading CGCs containing only one of these GH families would be excluded.

The phyla Bacteroidetes and Firmicutes contain the taxa with the highest numbers of mucin-degrading CGGs per genome (**Fig S7**). The genera *Bacteroides* and *Parabacteroides* contain the majority of strains with the highest number of mucin-degrading CGCs per genome (**Fig 5B**). Many of the strains that were enriched in humanized mice with PMG supplementation (unidentified *Bacteroides* spp., *B. ovatus, B. caccae*) were identified as strains in the HGM with the highest numbers of candidate mucin-degrading CGCs (**Fig 5B**). Proteobacteria, which were suppressed by PMG supplementation (**Fig 3C**) harbor very few mucin-degrading CGCs (**Fig S7**).

**Figure 5.**
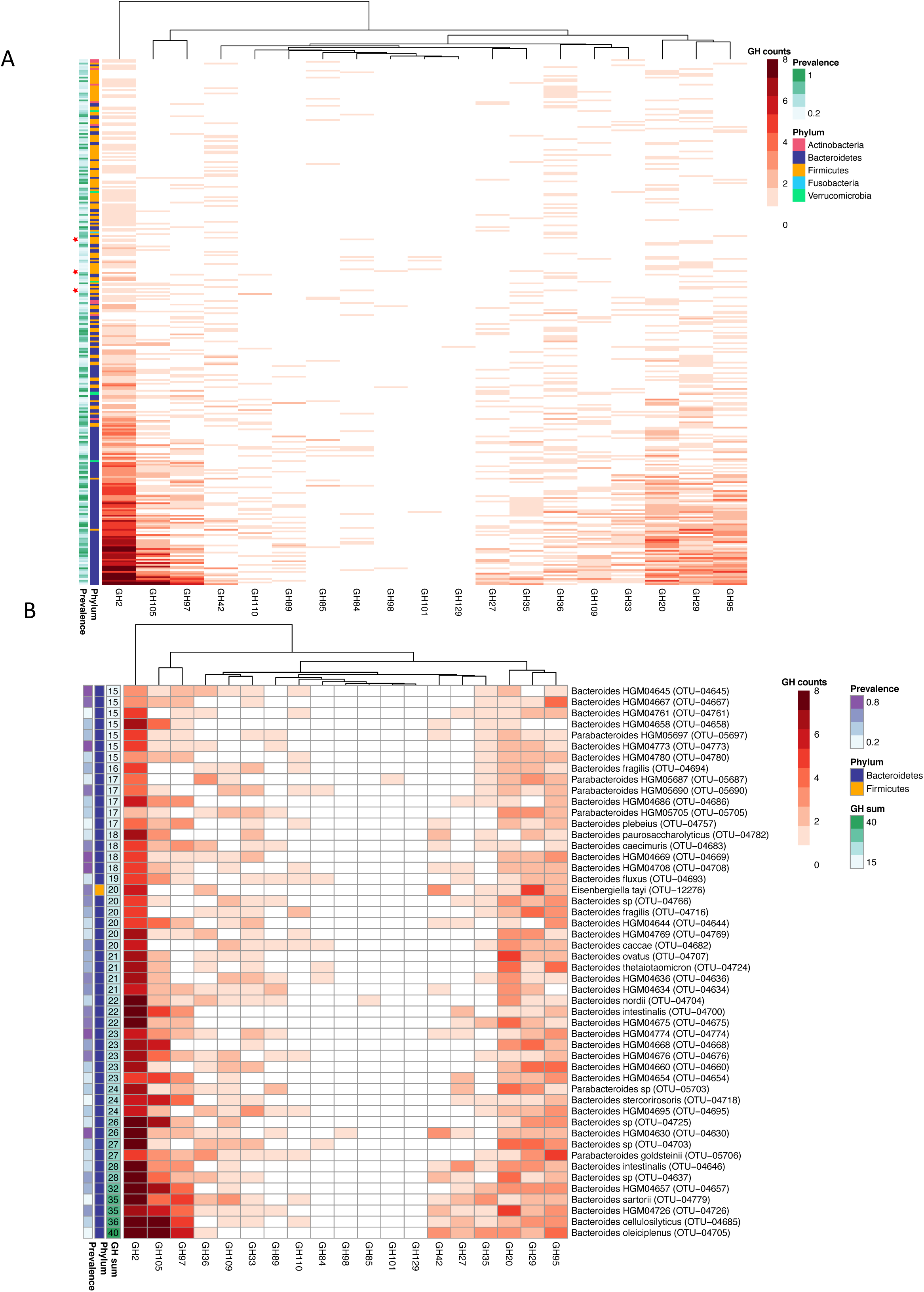
Prevalent gut commensals harbor high numbers of mucin-degrading carbohydrate gene clusters. **A)** Numbers of putative mucin-degrading CGCs per genome in 839 taxa that met criteria for at least 1 mucin-degrading CGC and 10% prevalence in healthy humans. A healthy cohort from the Human Microbiome Project (HMP) dataset is used to define prevalence. Stars indicate the pathogenic strains queried that contained putative mucin CGCs. **B)** The top 50 taxa within the HGM database harboring the highest number of mucin-degrading CGCs within their genomes. Total putative mucin-targeting glycoside hydrolases identified within these candidate mucin CGCs are indicated (GH sum).

Prevalent human commensal bacteria show a greater diversity and abundance of mucus-degrading CGCs than well-known enteric pathogens. Of the 34 species within the HGM database that we identified as enteric pathogens (please see Materials and Methods section and **Table S3**), only 4 met our criteria for harboring a candidate mucin CGC (*Vibrio vulnificus, Clostridium botulinum, Clostridium paraputrificum, Clostridium perfringens*), as opposed to 835 gut commensals. Of these, 299 strains are also prevalent (> 10%) in a healthy human cohort; only 3 are pathogens (**Fig 5A**). Taken together these results show that mucus utilization is enriched in commensal, gut-associated bacteria and not common enteric pathogens, consistent with mucus as a key component of the symbiosis between host and microbiota within the gut.

## Discussion

During the first months of life, delivery of glycans through breastmilk is an influential force that shapes the gut microbiota of newborns and infants. It has been suggested that HMOs are involved in healthy brain and body development[12, 42], immune system maturation[13, 16], and protection against pathogens[14, 15, 17, 18]. Only small amounts of intact HMOs are found in the feces and urine of breast-fed infants[43], suggesting that most HMOs are hydrolyzed by neonate gut commensals[44]. Our experiments comparing gnotobiotic and GF mice support this hypothesis, and our results demonstrate that the gut microbial community indeed metabolizes HMOs. Furthermore, selective consumption of exogenous glycans can influence the composition of microbes present in the gut, favoring microorganisms with a high diversity of carbohydrate-active enzymes (e.g., *Bt*[5]) or with particular hydrolase activity to cleave certain glycan linkages (e.g., *B. infantis*[22]). The metabolic flexibility of polysaccharide generalists such as *Bt* may confer a competitive advantage during the transition from infancy to adulthood, as it may shift its metabolism from HMOs to dietary or mucin glycans[24].

The pool of HMOs found in breastfeeding mothers varies in composition, structure and concentration day to day. When searching for molecules that aim to mimic the effect of HMOs on the gut microbiota, for example, in order to create an ideal infant feeding formula, similar structural complexity should be one of the major considerations. Our chromatographic mass-spectrometry analysis indicates that PMGs are a good candidate to mimic the complexity of HMOs. *In vivo* experiments with humanized mice indicate that PMGs and HMOs affect the microbiota in nuanced ways; the complex glycan mixtures structure the microbiota similarly at high-level ecological and taxonomic scales. A notably similar change due to supplementation with either glycan is an increase in the relative abundance of the species *A. muciniphila*, which relies on mucin as a carbon and nitrogen source, producing short chain fatty acids from mucin fermentation[45]. Several studies in humans and animal models have documented the presence of *Akkermansia* with various positive phenotypes for the host. Studies in pregnant women have shown *A. muciniphila*-like bacteria to be positively correlated with normal weight gain over pregnancy[46] and reduction of diabetes[38]. In addition, *Akkermansia* levels have been inversely related to the severity of obesity[47] and inflammatory bowel disease[48]. Colonization of obese mice with *A. muciniphila* reduces body weight without dietary change and reverse diet-induced fasting hyperglycemia and insulin resistance index, reducing adipose tissue[40]. Notably, recent studies have associated *A. muciniphila* with positive responses to immunotherapy in cancer patients[49] and maintenance of intestinal adaptive immune responses during homeostasis[50]. Studies demonstrating an enrichment of Verrucomicrobia in industrialized compared to traditional populations [51] and blooming in mice fed a fiber-poor diet coincident with inflammatory markers[1, 25] suggests that context, quantification of absolute abundance, and perhaps function and strain-specific features of *Akkermansia* may be important considerations in overall health impact of this taxon within an individual.

The broad spectrum of glycan structures present in breast milk may represent an ecological strategy in which a robust gut ecosystem is selected for via maternal transmission of microbes supported by HMOs. By delivering a diverse pool of glycans, the mother facilitates establishment of a microbial community that may be more stable than one which relies primarily on limited structural diversity (i.e., a single type of glycan) or on fluctuating dietary input from the host. The HMO-consuming community can utilize structurally similar mucins; we show here that bacteria prevalent in healthy humans possess a diverse array of glycoside hydrolases capable of mucin glycan degradation. The manner by which the complex chemistries of exogenously-administered PMGs influence the microbial community in the background of endogenous human mucus glycans versus murine mucus glycans is an important area of follow-up investigation. Previous work has demonstrated detrimental effects of microbial degradation of the host mucus layer[1, 25]; exogenously-provided mucus glycans may reduce the consumption of the host mucus in the absence of dietary MACs. The findings presented here support the idea that the host selects for a mucin-consuming microbial community to promote stability in the gut ecosystem. Accordingly, three diverse types of disturbance to the microbial community of humanized mice were mitigated by administration of exogenous PMGs. The close symbiotic interactions between commensals and their host appear to evolve at the mucus interface, with HMOs serving as important agents of this relationship after birth.

## Materials and Methods

### HMO isolation from human milk

Human milk was obtained from 12 healthy volunteers recruited at the UCSD Medical Center, San Diego, CA, after approval by the University’s Institute Review Board. Proteins and lipids were removed from milk samples with centrifugation and ethanol precipitation. Roto-evaporation was used to rid samples of residual ethanol. Lactose and salts were removed by gel filtration chromatography over a BioRad P2 column (100 cm x 16 mm, Bio-Rad, Hercules, CA. USA) using a semi-automated Fast Protein Liquid Chromatography (FPLC) system.

### HMO isolation and purification from animal specimens

HMOs were extracted from mouse ileum and feces, purified over C18 and Carbograph microcolumns, fluorescently labeled with 2-aminobenzamide (2AB) and separated by high performance liquid chromatography (HPLC) on an amide-80 column (4.6mm ID x 25cm, 5 μm, Tosoh Bioscience, Tokyo) with a 50 mM ammonium formate/acetonitrile buffer system. Separation was monitored by a fluorescence detector at 360 nm excitation and 425 nm emission. Peak annotation was based on standard retention times and mass spectrometric (MS) analysis on a Thermo LCQ Duo Ion trap mass spectrometer equipped with a Nano-ESI-source.

### HMO Profiling by High Performance Liquid Chromatography with Fluorescence Detection (HPLC-FL)

Isolated, dried HMOs from intestinal samples were fluorescently labeled with 2-aminobenzamide (2AB) and cleaned with silica spin columns as previously described. The 2AB-glycans were separated by HPLC-FL on an amide-80 column (4.6mm ID x 25cm, 5 mm, Tosoh Bioscience, Tokyo) with a linear gradient of a 50 mM ammonium formate/acetonitrile buffer system. Separation was performed at 25° C and monitored by a fluorescence detector at 360 nm excitation and 425 nm emission.

### Competitive colonization of gnotobiotic mouse

Germ-free Swiss-Webster mice were reared in gnotobiotic isolators and fed an autoclaved polysaccharide-deficient diet (BioServ, http://bio-serv.com) in accordance with A-PLAC, the Stanford IACUC. Mice were bi-colonized with overnight cultures of *Bt* and *B. infantis* using oral gavage as described in [5]. Subsequent community enumerations from mice were determined from freshly collected feces, by selective plating of serial dilutions on Reinforced Clostridial Media (RCM) agar and Brain-Heart Infusion (BHI)-blood agar supplemented with gentamicin (200 µg/ml). Significant differences between sample groups were determined using Student’s t-test.

### LNT and LNnT synthesis

Synthetic LNT and LNnT (Glycom A/S) were crystallized to a final purity of >99%. Characterization was performed using multiple methods including NMR (1D and 2D) mass-spectrometry, and HPLC.

### PMGs preparation

PMGs were prepared as described by Martens *et al*[30], with some modifications: *O*-glycans were released from porcine gastric mucin (Sigma Type III, 10% w/v) by incubation at 48**°**C for 20 hrs in 150 mM NaOH with 750 mM NaBH_4_. The reaction was neutralized with HCl (10 M). Insoluble material was removed by centrifugation at 14000 x *g* (30 mins, 4° C). Supernatant was filtered and dialyzed against dH_2_O with 1kD MWCO membranes (Spectra/Por 7, Spectrum Labs)) and subsequently lyophilized. Glycans were solubilized in 50mM Tris pH 7.4 buffer and fractionated using DEAE-Sepharose CL-6B anion exchange columns.

### Monosaccharide and sialic acid determination from PMGs

For monosaccharide composition analysis, PMG samples were hydrolyzed using 2 M trifluoroacetic acid (TFA) at 100°C for 4 h followed by removal of the acid under dry nitrogen flush. Dried samples were co-evaporated with 50 μl aqueous iso-propyl solution (50% IPA) twice to ensure complete removal of TFA. Finally, samples were dissolved in water and analyzed by HPAEC-PAD (Dionex ICS3000) using a CarboPac PA-1 (4×250mm0 column with 100mM NaOH and 250mM NaOAc. Monosaccharides were identified with a sensitive post-acceleration detector (PAD) using standard Quad potential as specified by the manufacturer.

Sialic acid content of PMGs was determined after hydrolysis with 2M HOAc at 80° C for 3 hours. Acetic acid was removed by speed vacuum and free sialic acids collected by spin-filtration through a 3k MWCO filter followed by derivatization with 1,2-diamino-4,5-methyleneoxybenzene (DMB). Fluorescently-labeled sialic acids were analyzed by reverse-phase HPLC using Acclaim120 C18 column (4 × 250mm, 5µ, Dionex) and a fluorescence detector with a slow gradient of 9% to 14% acetonitrile over 20 min.

### MALDI-mass spectrometry method for PMG structural analysis

Samples were prepared for MALDI-mass spectrometry with slight modifications to an established method[52]. To per-*O*-methylate, PMG samples were dissolved in dry DMSO, to which 100 µl sodium hydroxide in DMSO was added, followed by immediate addition of 200 μL methyl iodide. After continuous stirring for 1 h, a second aliquot of methyl iodide (50 μL) was added, followed by stirring for 30 min. The reaction was stopped with ice-cold water and the permethylated glycans were extracted with 1 mL chloroform. The chloroform layer was washed twice with water, dried, re-suspended in methanol, mixed with sDHB matrix and spotted on a MALDI plate. Spectra were acquired on positive mode, and putative glycan structures were assigned with Glyco Workbench[53, 54].

### Mouse humanization and enumeration

Germ-free Swiss-Webster mice were humanized once with 200 µl of frozen human fecal sample resuspended 1:1 in anaerobic phosphate buffer saline (PBS) by oral gavage. The same adult human donor sample was used for all humanization experiments. The humanized microbiota was allowed to equilibrate for 4-6 weeks prior to onset of experimental manipulation. Mice were maintained in gnotobiotic isolators throughout the duration of the experiments and fed one of three diets: standard (Purina LabDiet 5K67), polysaccharide-deficient (BioServ, http://bio-serv.com) or high fat (60% fat and 20% carbohydrates, D12492, Research Diets Inc.). When needed, 1 mg of clindamycin (Sigma) per mouse was administered via oral gavage or 200 µl of an overnight culture of *C. difficile* growth in RCM (BD Difco). To approximate the mass of HMOs available to breastfeeding infants, complex pools of glycans (purified HMOs and PMGs) were added at a concentration of 1% (w/v) in water unless otherwise specified. Infants consume approximately 7.5-11.25 g HMOs daily (∼750 mL milk intake, HMOs are 10-15 mg/mL in breastmilk), 1.5-2.25 g/kg (for a 5 kg infant). Mice consume about 5 mL water per day; at 10 mg/mL glycan concentration, this equates to 2 g/kg for a 25 g mouse. LNT and LNnT together constitute 15-20% total HMOs in breastmilk[8]; as such, administration of pure glycans (LNT, LNnT) are provided at a higher concentration than at which they naturally occur. All animal protocols were in accordance with A-PLAC, the Stanford IACUC.

### C. difficile *enumeration*

For quantification of *C. difficile* CFU, 1 μl feces was serially diluted in PBS and plated onto selective media, composed of Clostridium difficile Agar Base (OxoiD) with 7% v/v of Defibrinated Horse Blood (Lampire Biological Laboratories), supplemented with 32 mg/L Moxalactam (Santa Cruz Biotechnology) and 12 mg/L Norfloxacin (Sigma-Aldrich) (CDMN). Plates were incubated overnight at 37° C in an anaerobic chamber (Coy). Identification of colonies as *C. difficile* was validated by colony PCR, using the primers Cl1 (5’-TGTTGCAATATTGGATGCTTT) and Cl2 (5’-TGACCTCCAATCCAAACAAA), which target a fragment of *tcdB* gene.

### 16S rRNA amplicon sequencing and analysis

Fresh fecal samples were collected and frozen at -80° C. DNA was extracted according to Earth Microbiome Project standard protocols using the Powersoil-htp extraction kit (MoBio). The 16S rRNA gene was amplified (515F, 806R) and sequenced using the Illumina MiSeq platform at the Medical Genome Facility, Mayo Clinic, Rochester, MN across 6 runs at 150 bp except the experiments with HFD, where 300 bp (Figures 4A,B,E,F,G) and 250 bp (Figures 4C,4D,S6) were generated. Raw reads were demultiplexed using QIIME 1.9[55] and subsequently trimmed and denoised using DADA2 with standard input parameters maxN=0, maxEE=2, trunQ=2 except for the GOS/HMO/PMG supplementation experiment (corresponding to Figures 1C,1D,S3) where maxEE=(5,2) was used[56]. Taxonomy was assigned with the GreenGenes training set version 13.8 clustered at 97% identity. Phylogenetic trees were constructed by performing a multiple sequence alignment and constructing a Generalized time-reversible with Gamma rate variation maximum likelihood tree using a neighbor-joining tree as a starting point with the R packages *msa* and *phangorn* as described previously[57]. All resulting datasets were filtered for low-abundance ASVs at 10% prevalence and subsequently rarefied. R packages *phyloseq, DESeq2, RandomForests, rfUtilities, vegan, caret, cluster, harrietr, ggpubr, plotrix, rstatix* and *ggplot2* were used for normalization, analyses, and visualization.

### *Mining for mucin carbohydgrate gene clusters* in silico

Four thousand five hundred and fifty eight fully sequenced genomes were selected based on their identification as being human-associated (n=4558, in the Human Gut MAG Species Database, HGMdb; https://github.com/snayfach/IGGdb)[41]. Carbohydrate-active enzyme assignment of glycoside hydrolases (GHs) was performed using a Hidden Markov Model database[58, 59]. We identified physically linked carbohydrate gene clusters (CGCs) using a default stringent parameter in the dbCaN software, which defines a CGC as physical linkage (distance ≤ 2) of at least two CAZymes with a transcription factor (TF) or transporter (TC); in this case CAZymes were constrained to those listed in **Table S2**. GHs within CGCs were summed across each genome, and heat maps were generated using CGC counts. Prevalence of taxa across individuals was determined by running the IGGSearch software [41] with default settings on the 180 metagenomes available from healthy individuals in the HMP (https://hmpdacc.org/). Taxa were defined as pathogenic if they met the criteria of belonging to a list of genera (*Salmonella, Shigella, Yersinia, Vibrio, Campylobacter*) or species (*Clostridium difficile, Clostridium perfringens, Clostridium botulinum, Pseudomonas Aeruginosa*), and could be manually verified as pathogens based on existing literature (see **Table S3** for a list of these 34 species).

### Body composition analysis

Animals were measured for total body fat mass and adipose deposits were precisely dissected and weighted. Short chain fatty acid content was determined as described previously[60].

### Data Availability

All raw 16S sequencing data will be made publicly available. Code will be made available upon request.

## Acknowledgements

We thank members of the Sonnenburg Lab for helpful discussions. We thank the UCSD Glycotechnology Core Facility for technical assistance and Glycom for their generous contribution of LNT and LNnT. Evelyn Jantscher-Krenn and Alex Szyszka at the Division of Neonatology and Division of Gastroenterology and Nutrition, Department of Pediatrics, UCSD, provided technical support for the HMO isolation. The authors acknowledge support from the National Institutes of Health R01-DK085025 (to J.L.S.). J.L.S. is a Chan Zuckerberg Biohub Investigator.

## Figure Legends

**Figure S1.**
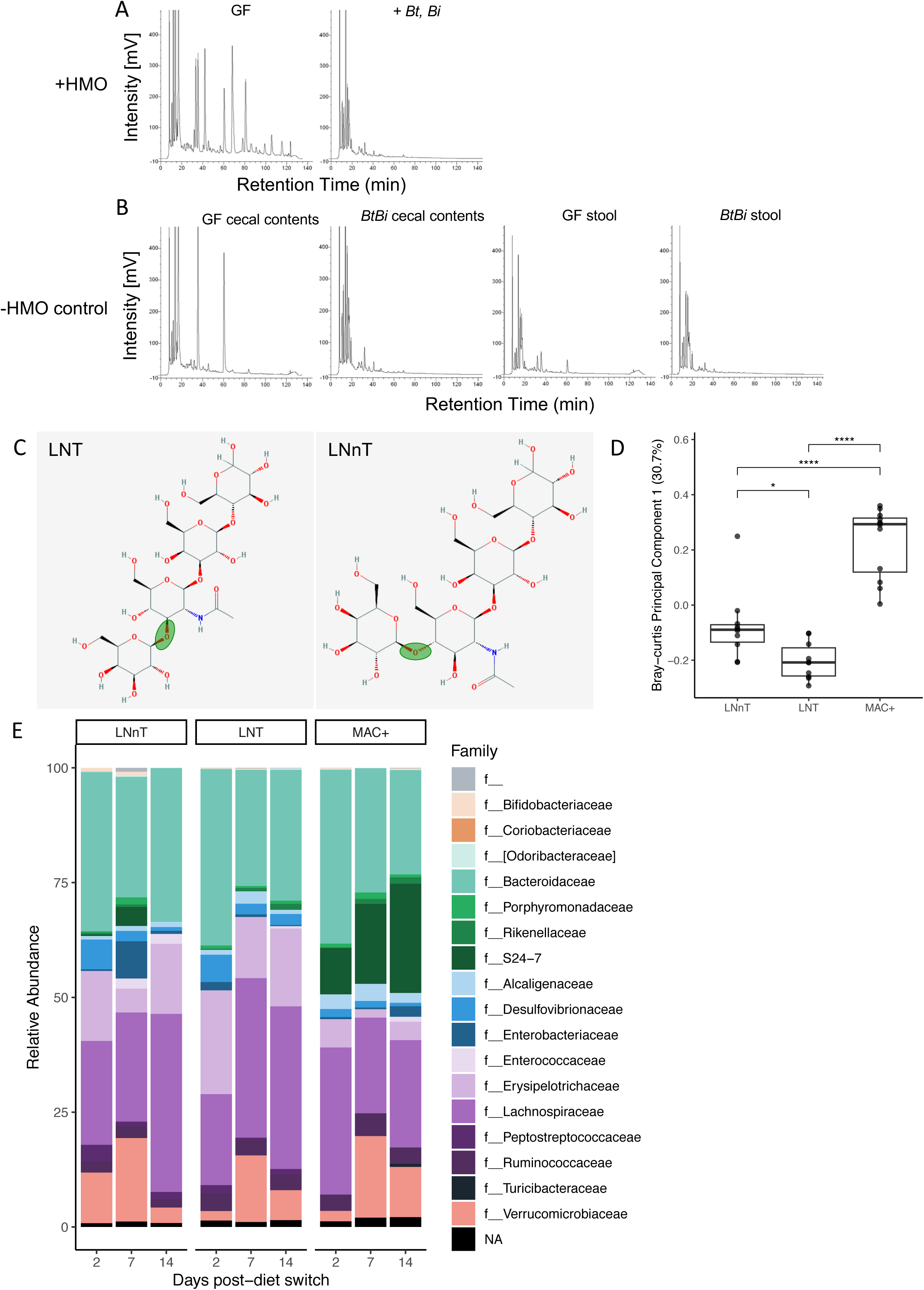
**Corresponds to Figure 1. A)** HPLC-FL based chromatograms of glycan content in cecal contents of germ-free or bi-colonized mice fed 1% HMOs in drinking water. At day 7, mice bi-colonized with *Bt* and *B. infantis* degrade the HMOs still visibly present in germ-free mice. **B)** Glycan content from germ-free and bi-colonized control (*BtBi* = *B. theta, B. infantis*) mice not fed HMOs. **C-E)** Supplementation with two synthetic HMOs that differ in a single glycosidic linkage affects the composition of the gut microbiota. **C)** Structure of LNT and LNnT, two synthetic HMOs. Green circles indicate the differing location of the Gal-Glc linkage. **D)** Bray-Curtis distance reveals full separation of microbial communities of mice on MAC^+^ diet from MD supplemented with either synthetic HMO, as well as separation of LNT from LNnT (F_(2,29)_=41.736 * *P* < 0.05, **** *P* < 0.0001, one-way ANOVA with Tukey’s post-hoc comparisons). **E)** Relative abundance of bacterial families in MAC^+^ or MD supplemented with LNnT or LNT. f_ indicates that a strain is not assigned at the family level in the Greengenes database; NA indicates lack of taxonomic assignment at the family level.

**Figure S2.**
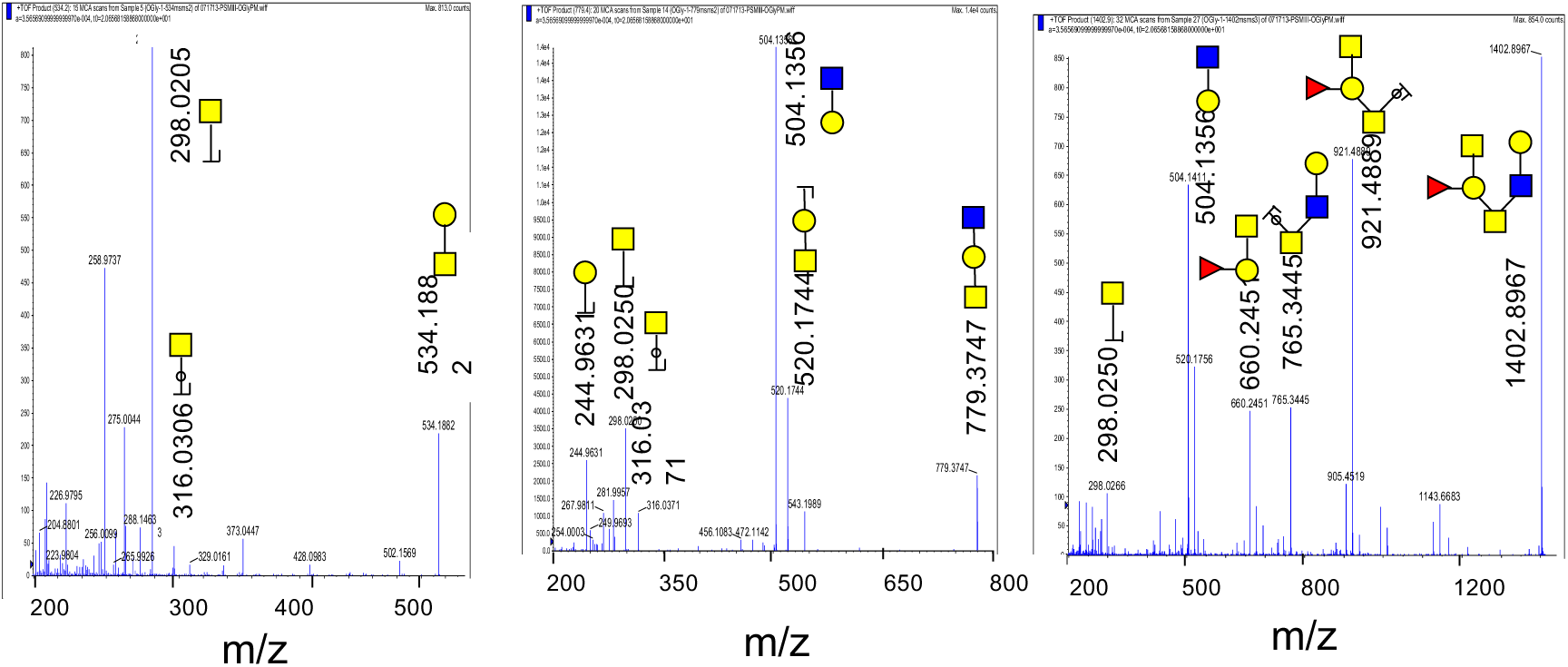
**Corresponds to Figure 2.** Verification with MS/MS fragmentation pattern of the three porcine mucin glycan structures predicted by GlycoWork Bench from Fig 2C.

**Figure S3.**
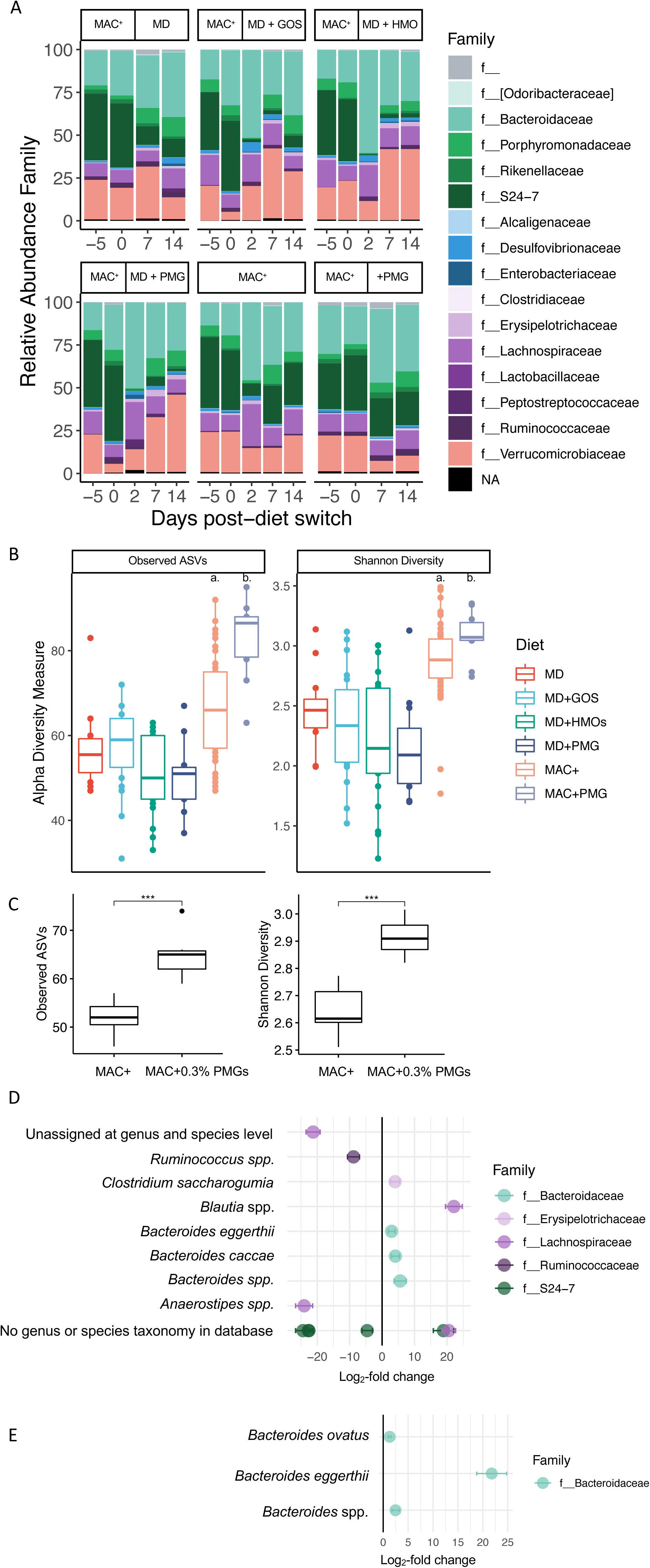
**Corresponds to Figure 1. A)**. Diet-induced changes to the relative abundance of bacterial families over time. All mice began on MAC^+^ baseline diet days prior to glycan supplementation. f_ indicates that a strain is not assigned at the family level in the Greengenes database; NA indicates lack of taxonomic assignment at the family level. **B)** There are no differences in two alpha diversity metrics between groups on MD dietary background; however, 1% PMG supplementation to MAC^+^ diet results in significant alpha diversity enrichment compared to MAC^+^ alone. Observed ASVs and Shannon index are higher in MAC^+^ +/- PMGs than MD supplemented with various glycans (observed ASVs: F_(5,128)_=17.382, *P* < 0.0001 one-way ANOVA with tukey’s post-hoc comparisons. a. * MAC^+^ vs. MD+GOS, **** MAC^+^ vs MD+HMOs, *** MAC^+^ vs. MD+PMG, *** MAC^+^ vs MAC^+^+PMG. Shannon diversity: F_(5,34.74)_=20.83, *P* < 0.0001 Welch’s ANOVA with Games-Howell post-hoc comparisons. b, **** MAC^+^+PMG vs. MD, MD+GOS, MD+HMOs, and MD+PMG.) **C)** More dilute PMG supplementation (0.3%) still leads to significant enrichment of alpha diversity to MAC^+^ diet (*** *P* < 0.001, Student’s t-test). **D)** 1% PMG supplementation (positive log_2_-fold change) to MD diet (negative log_2_-fold change) led to significant enrichment of several taxa that were also enriched by HMOs (mean +/- SEM, Wald Test, adjusted *P* value < 0.05). **E)** 0.3% PMG supplementation to MAC^+^ diet (as in S3C) led to enrichment of three *Bacteroides* species (mean +/- SEM, Wald Test, adjusted *P* value < 0.01).

**Figure S4.**
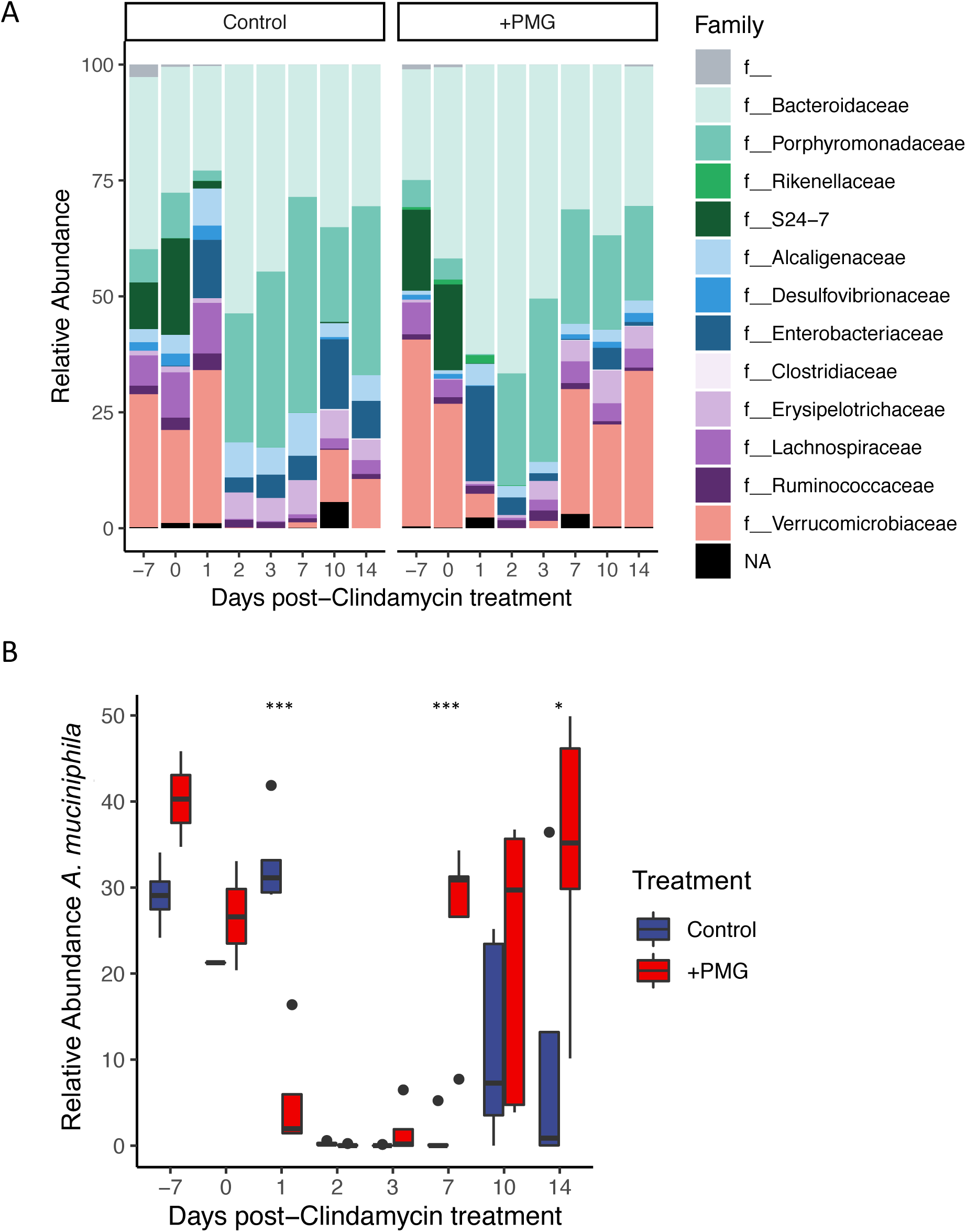
**Corresponds to Figure 3. A)** Relative abundance of bacterial families after clindamycin treatment. f_ indicates that a strain is not assigned at the family level in the Greengenes database; NA indicates lack of taxonomic assignment at the family level. **B**. PMGs supplemented to MD diet lead to enhanced recovery of *A. muciniphila* (Pairwise t-tests with Bonferroni correction, * *P* < 0.05, *** *P* < 0.001).

**Figure S5.**
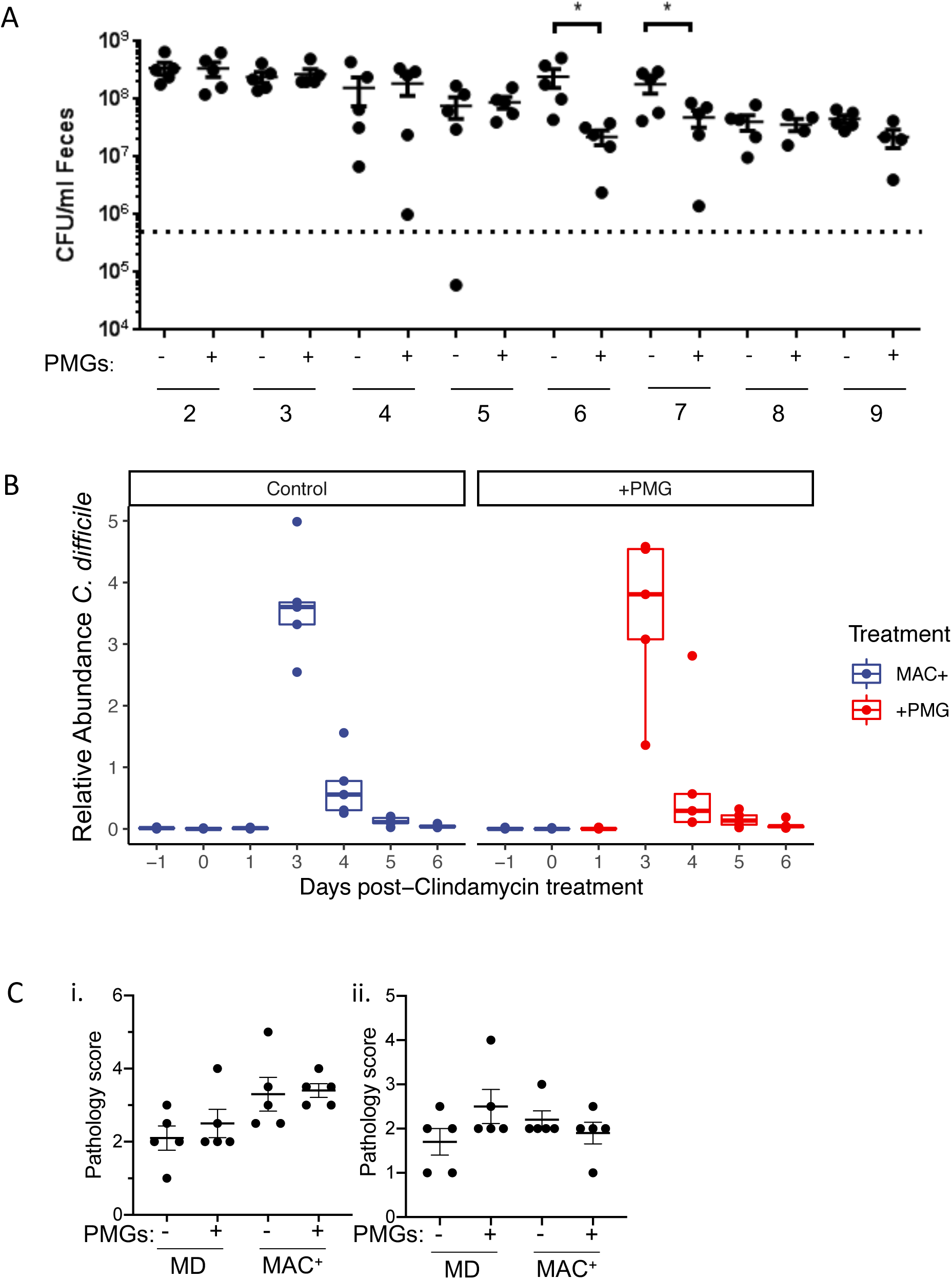
**Corresponds to Figure 3. A)** Absolute abundance of *Cd* colonization enumerated with selective plating is reduced with PMG administration days 6 and 7 post-infection compared to unsupplemented control (MD diet, * *P* < 0.01, Student’s t-test). **B)** Relative abundance of *Cd* is not affected by PMGs in a MAC^+^ background. **C)** Blinded histopathological scoring of cecal (**i**.) and distal colon (**ii**.) tissues from mice infected with *Cd*.

**Figure S6.**
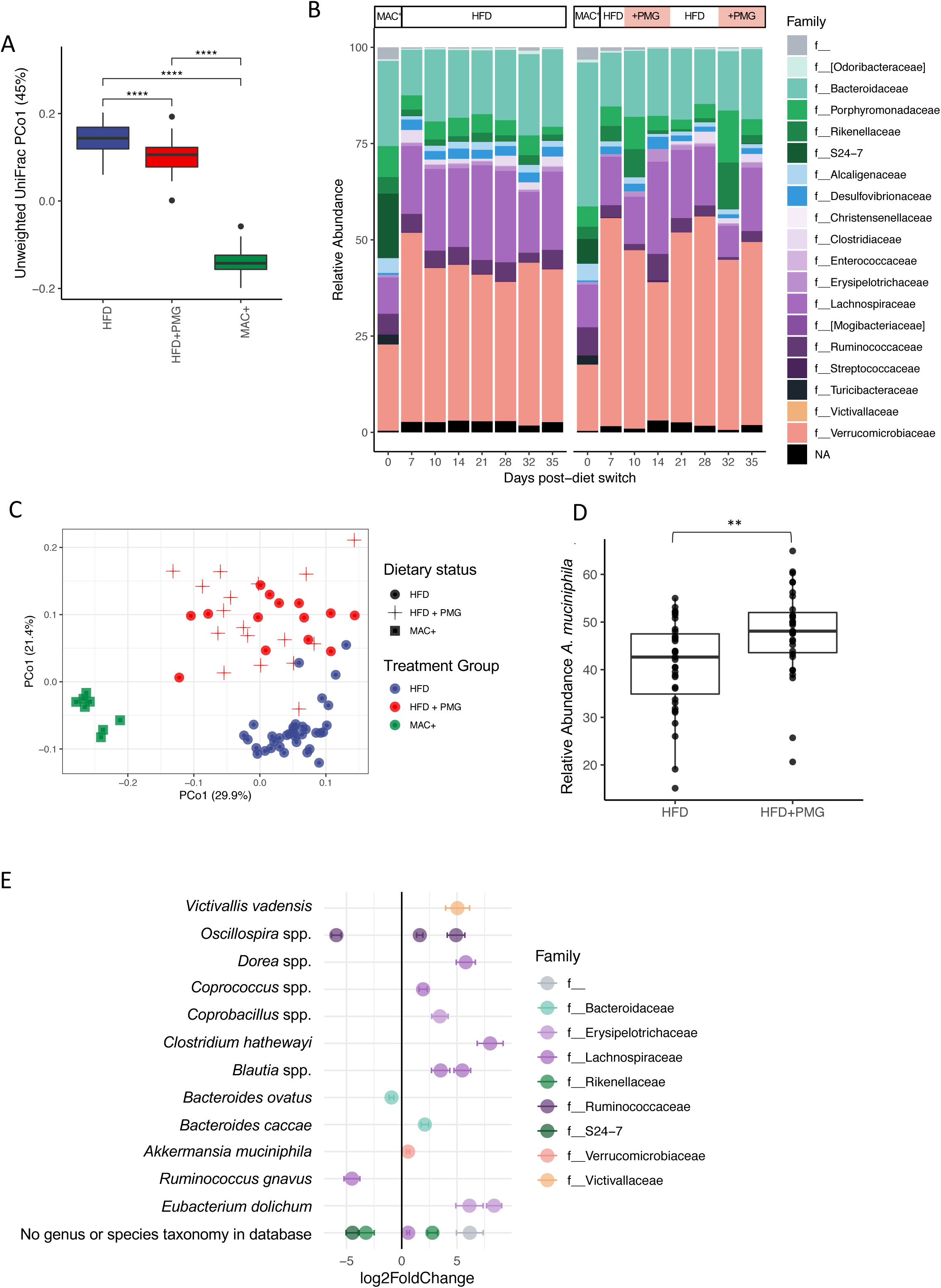
**Corresponds to Figure 4. A)** PMG supplementation to HFD leads to distinct communities from HFD alone or MAC^+^ diet as quantified by the first principal component of unweighted UniFrac distance between communities (F_(2,107)_=782.603 **** P < 0.0001 ANOVA with Tukey’s post-hoc comparisons. n=30 HFD, n=29 HFD+PMG, n=51 MAC^+^). **B)** Relative abundance of the top 100 most abundant taxa on HFD alone or HFD with transient (1 week) PMG supplementation. Salmon-colored background boxes indicate sampling timepoints during which 1% PMGs were administered to the latter group. Both groups started on MAC^+^ diet (day 0). f_ indicates that a strain is not assigned at the family level in the Greengenes database; NA indicates lack of taxonomic assignment at the family level. **C)** Microbial communities of mice that were treated transiently with PMGs (red crosses) remain distinct from mice on HFD alone (blue), even when PMGs are removed from HFD (red circles). **D)** *A. muciniphila* reaches a higher relative abundance in mice treated transiently with PMGs (** *P* < 0.01, t-test). **E)** Taxa that are significantly enriched due to transient PMG supplementation (positive) in water to HFD background (negative, mean +/- SEM, adjusted *P* value < 0.0005).

**Figure S7.**
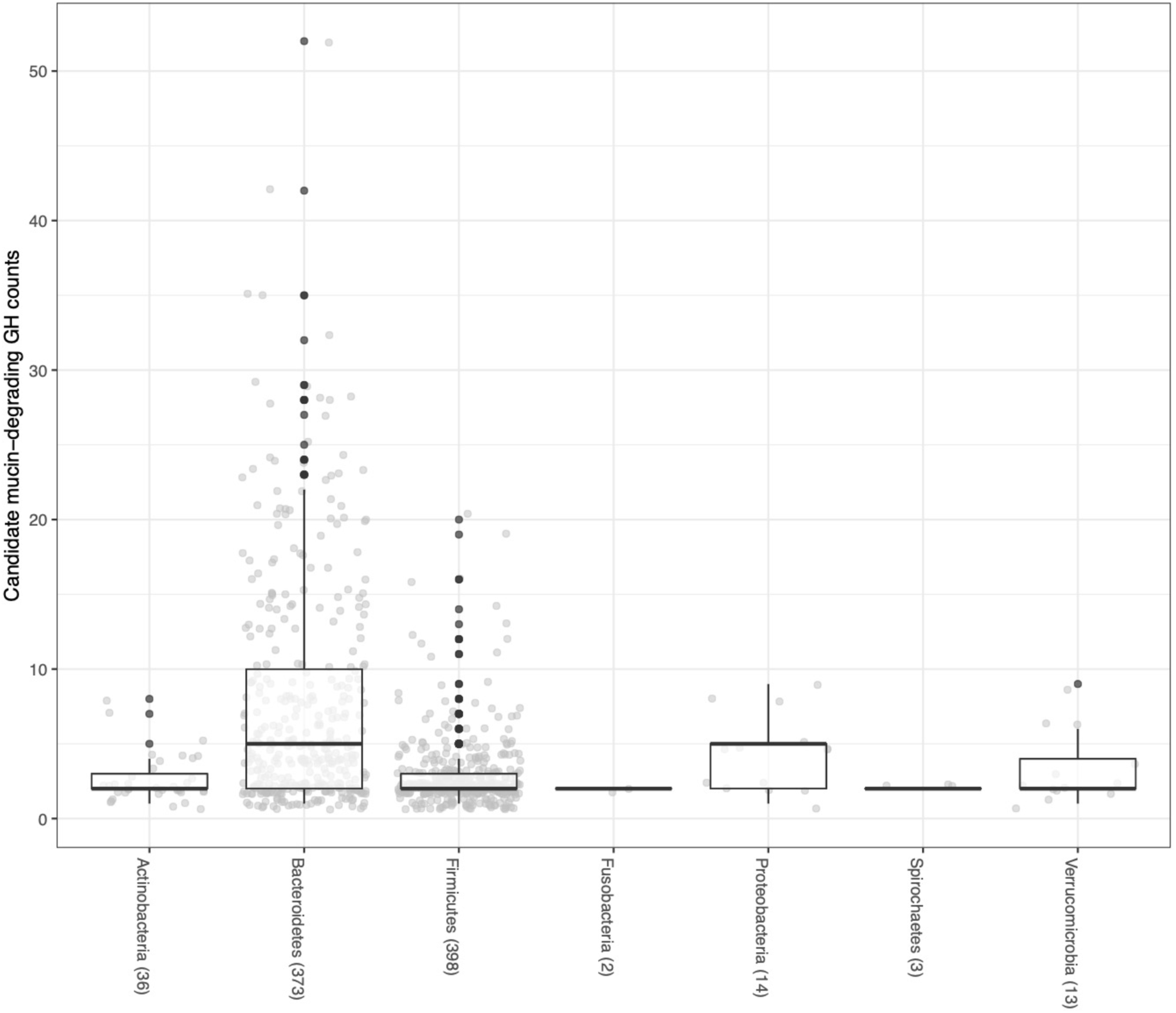
**Corresponds to Figure 5. A)** The distribution of number of putative mucin-degrading CGCs per genome amongst all phyla in the HGM database.

**Table S1.**Top 20 predictive taxa of the Random Forests classifier to predict HFD or HFD with transient PMG supplementation (Figure 4C,4D,S6B-E.).

**Table S2.**List of literature references for mucin-targeting GHs[25, 61–64].

Table S3.List of bacteria categorized as pathogens for the mucin-glycan degradation analysis presented in Figure 5.

